# Meta-analysis reveals that sexual signalling in animals is honest and resource-based

**DOI:** 10.1101/2020.08.21.261081

**Authors:** Liam R. Dougherty

**Affiliations:** Department of Evolution, Ecology and Behaviour, University of Liverpool, Liverpool, L69 7RB, UK, Email

**Keywords:** Sexual signalling, courtship, condition-dependence, terminal investment, lek, handicap, systematic review

## Abstract

Animals often need to signal to attract mates, and sexual signalling may impose significant energetic and fitness costs to signallers. Consequently, individuals should strategically adjust signalling effort in order to maximise the fitness payoffs of signalling. An important determinant of these payoffs is individual state, which can influence the resources available to signallers, the likelihood of mating, and the motivation to mate. However, empirical studies often find contradictory patterns of state-based signalling. For example, some studies find that individuals in poor condition signal less, in order to conserve resources (ability-based signalling). In other cases, individuals in poor condition signal more, in order to maximise short-term reproductive success (needs-based signalling). I used meta-analysis to examine animal sexual signalling behaviour in relation to six aspects of individual state: age, mated status, attractiveness, body size, condition, and parasite load. Across 228 studies and 147 species, individuals (who were overwhelmingly male) signalled significantly more when in good condition, and there was a strong positive trend for increased signalling for large, attractive individuals with a low parasite load. Overall, this suggests that animal sexual signalling behaviour is generally honest and ability-based. However, needs-based signalling (terminal investment) was found when considering age, with old virgins signalling more than young virgins. Sexual signalling was not significantly influenced by mated status. There was a large amount of heterogeneity across studies that remained unexplained, and therefore more work is needed to determine the ecological factors influencing the magnitude and direction of state-dependent sexual signalling.

## Introduction

Sexually-reproducing animals often need to signal to attract mates^1^. Signalling can be morphological or behavioural, and individuals may signal over a long range in order to attract potential partners (advertising their presence), and over a short range in order to convince potential partners to mate (advertising their quality as a mate; short range signalling is often referred to as courtship). However, signalling is often costly. For example, behavioural displays require energy, and though the energetic cost of a single display bout may be small^2^, the long term costs may be large if displays are repeated often and over a long time^3,4^. Further, as well as direct energetic costs, sexual signalling may also impose indirect fitness costs. For example, engaging in long signalling bouts may reduce survival^5^, and both morphological and behavioural signals can increase the conspicuousness of signallers to predators^6,7^. These costs are exacerbated by the process of sexual selection, whereby the mating preferences of choosers can drive the evolution of larger and more elaborate display traits and behaviours in courters^1, 8^.

Given that signalling is often costly, the resources available to an individual should influence the amount of signalling they can afford to do. In other words, signalling may often be condition-dependent^9,10^. For example, poor-condition individuals often have smaller or less elaborate ornaments than high-condition individuals^9^. Indeed, one theory to explain the evolution of elaborate ornaments is that they honestly signal individual condition or quality: when ornaments are costly to produce, they are hard to fake^11,12^, because of the inherent energetic trade-offs between investment in reproductive versus somatic processes. Behavioural displays may reflect individual quality or condition, especially if they are elaborate or difficult to perform^2,13,14^. Poor-condition individuals may therefore show reduced signalling behaviour, in terms of time spent displaying, or the complexity or energy content of a display, because they have limited energy reserves. This can be considered an ‘honest’, or ‘ability-based’ strategy (*sensu* ^15^). Importantly, the amount of resources individuals are able to allocate to signalling could depend on several intrinsic factors, including age, body size, parasite load, and stress levels^9,16,17^.

A key difference between morphological and behavioural signals is that the latter are inherently dynamic: behaviour can vary over very short timescales. This means individuals could strategically vary their signalling behaviour in order to maximise the fitness payoffs of signalling^16^. For example, signallers could benefit from only signalling when predation risk is low^6^ (though recent work suggests that animals do not consistently alter their signalling behaviour in response to the environment^18^). Alternatively, individual condition or state may influence the fitness payoffs of signalling. For example, virgins benefit more from mating than once-mated individuals, and so, all else being equal, virgins should invest more into signalling in order to try and achieve a mating^19,20^. Additionally, if individuals in poor condition are less attractive to potential mates, then a given courtship bout is less likely to result in successful mating^21,22^. Therefore, rather than court uninterested partners, poor-condition individuals could instead choose to forego signalling and conserve energy until their condition improves. Taken together, this means that a reduction in signalling behaviour for poor condition individuals could arise for two reasons: either because they have fewer resources available to allocate to signalling, or as an adaptive strategy in response to current or expected fitness payoffs. Alternatively, old or poor-condition individuals could instead elect to *increase* their investment in signalling (and other reproductive traits) in order to try to maximise their short-term reproductive success at the expense of survival^23^. This can be considered a ‘dishonest’, or ‘needs-based’ strategy (*sensu* ^15^), and is often referred to as ‘terminal investment’ when applied to age^24,25^. Maximising short-term reproductive success will clearly benefit old individuals, who may have few remaining chances to mate. It may also benefit poor-condition individuals (of all ages) if they have an increased mortality risk, or a decreased mate encounter rate, compared to high-condition individuals^25^.

Empirical evidence for state-dependent plasticity in sexual signalling is widespread in animals, in relation to a range of intrinsic factors. However, for many factors there is mixed or conflicting evidence in relation to the *direction* of plasticity. For example, signalling has been shown to be higher in both good-condition^26,27^ and poor-condition individuals^23,28^, in both young^29,30^ and old individuals^31,32^, and in both large^22,27^ and small individuals^21,33^. Further, other studies have found no significant effect of these factors on signalling behaviour^34-36^. This variation could reflect the fact that different species show different state-dependent strategies, perhaps moderated by additional biological or environmental factors. For example, a strategy maximising short-term reproductive success might be more common for short-lived species, members of which cannot typically afford to wait for their circumstances to change in later life. Nevertheless, in cases such as this, where empirical results are mixed across a wide range of studies, there is a danger in simply cherry-picking confirmatory studies, or simply counting significant and non-significant results^37^. Instead, meta-analysis, taking into account the size and direction of effects (rather than simply P values), can be used to estimate the overall evidence for state-dependent plasticity in signalling, whilst also taking account of potential moderating factors that contribute to variation in plasticity^37^.

I investigated the extent to which sexual signalling behaviour is state-dependent in animals using a meta-analysis approach, by first systematically searching the published literature for estimates of between-individual variation in signalling behaviour. I focused on six state factors which may influence either the resources available to signallers, the motivation to mate, or the expected fitness payoffs associated with sexual signalling: age, attractiveness, body size, condition, mated status (whether individuals are virgin or mated), and parasite load. For all of these factors (with the exception of mated status), there is theory to predict plasticity in signalling in either direction, depending on whether energetic resources or the potential fitness payoffs of signalling are more important (see Factors section). I use this analysis to ask three questions. First, does individual state significantly influence signalling behaviour across the animal kingdom? Second, does signalling behaviour generally reflect the resources available to the individual (honest, ability-based signalling) or the fitness payoffs associated with signalling (needs-based signalling)? Third, are there any biological or methodological factors which influence the magnitude or direction of plasticity?

## Methods

### Literature searches

As part of a broader project, I searched for studies examining state-dependent variation in three mating behaviours: sexual signalling, response to sexual signals (responsiveness) and the strength of mate choice (choosiness). However, I only report the results for sexual signalling here (the responsiveness and choosiness results will be presented in a forthcoming paper). I searched for relevant papers in two ways. First, I obtained all papers cited by two reviews of state-dependent mating behaviour (with a focus on mate choice): Cotton *et al*.^38^ and Ah-King & Gowaty^39^. I also searched the online databases Web of Science and Scopus for all studies citing these two reviews on the 13/08/2019. Second, I performed literature searches using the online databases Web of Science & Scopus on the 13/08/2019. Screening of abstracts was performed using the Rayyan website^40^. The search and study selection process is outlined in **Figure 1**. Full search terms and further details are available in the supplementary methods.

**Figure 1.**
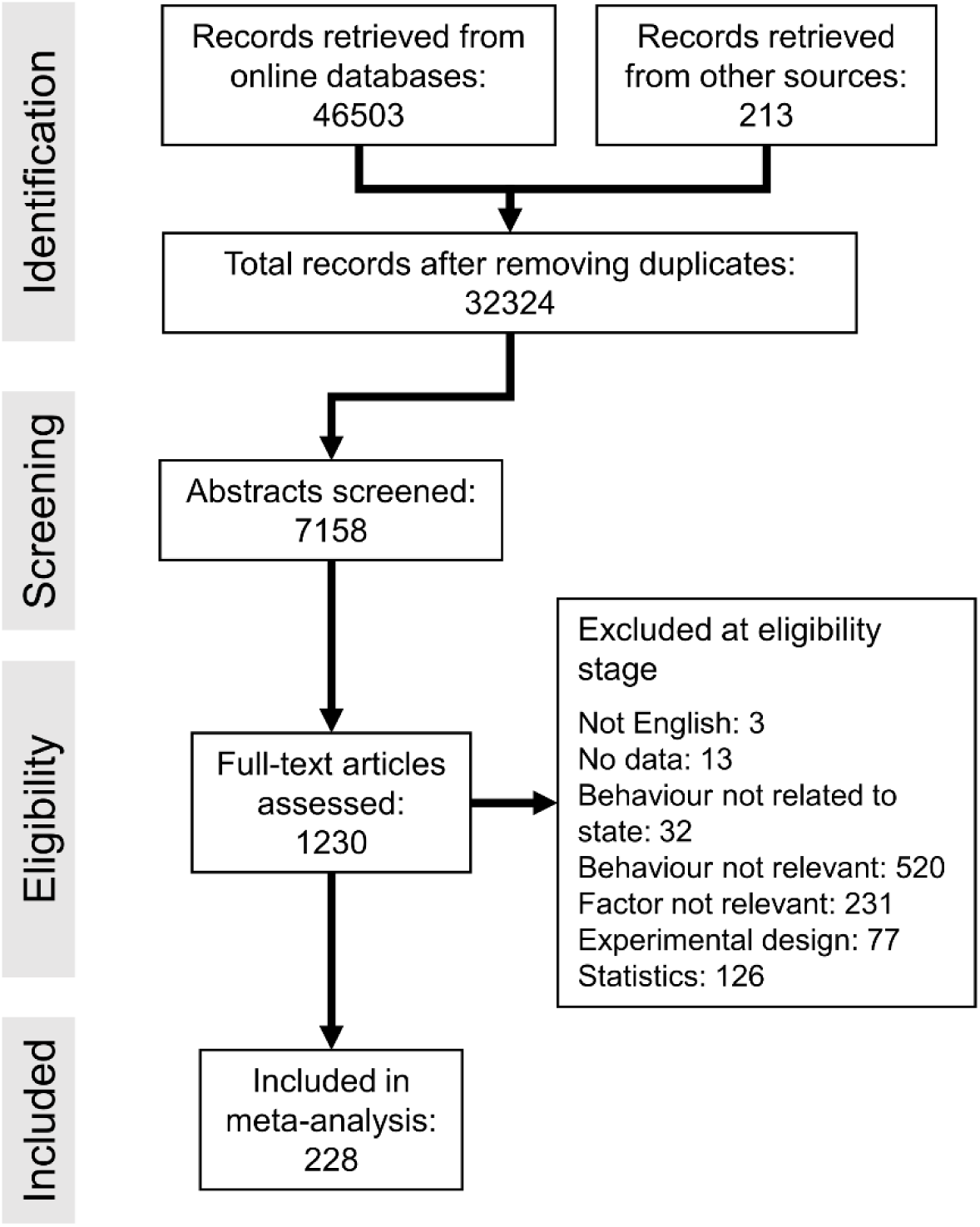
PRISMA diagram showing the literature search and screening process. Note that my search terms included a broader range of behaviours than presented in this study (see text for details).

### Criteria for study inclusion

A study was considered relevant if it linked between-individual variation in animal sexual signalling (excluding humans) to one of six state factors (age, attractiveness, body size, condition, mated status, or parasite load), and presented appropriate statistics or raw data so that I was able to calculate an effect size. I focused on sexual signalling behaviour, including long-range attraction signals and short-range courtship signals, but excluded less dynamic morphological signals such as ornaments or advertising colours. I excluded intrasexual signals, or signals for which a primary intrasexual function could not be ruled out. However, I acknowledge that all sexual signals probably function intra-sexually to some extent. I also included several lekking species for which displays probably signal to potential mates and rivals, because I consider the primary function of leks to be mate assessment. I included studies examining both male and female sexual signalling.

In terms of signalling behaviour, I included acoustic, visual, olfactory (pheromone) and tactile signalling. I focused on behavioural traits that reflect motivation to signal (courtship latency), or energetic investment in signalling (such as signalling duration, rate, and some measures of intensity). For acoustic signals, I included measures of call loudness except when related to body size (because call loudness may be constrained by the size of the sound-producing organs). I excluded measures of signal complexity, because this does not necessarily relate to overall energetic investment *per se*. For acoustic signals, I excluded measurements of call pitch/frequency and fine-scale temporal components of a call. For pheromones, I excluded measurements of pheromone composition, but included measures of time spent releasing pheromone, and pheromone titre if measured outside the body (I excluded measures of pheromone titre in dissected glands or bodies).

### State factors

I focused on six aspects of individual state which could influence sexual signalling. See the supplementary background for a more detailed discussion of the ways in which these factors may influence signalling behaviour.

#### Age

I only considered age-dependent effects if all individuals were sexually-mature, and age was not confounded with body size. Importantly, age is often confounded by mated status, especially for wild individuals, and this may influence the motivation to signal independently of age. Therefore, I only considered studies examining age-related signalling in virgins. This was necessary because few studies record both age and mated status in a way that allows their independent effects to be estimated.

#### Attractiveness

I assigned studies measuring several behavioural and morphological traits to this category, if the focal trait was hypothesised to signal mate quality (either genetic quality or current condition), or has been shown to be used in mate choice. I included studies measuring: a) song quality, b) ornament size, c) ornament or body colouration or brightness, d) morphological asymmetry, e) inbreeding, f) territory or nest quality and g) social rank.

Individuals were assumed to be attractive if they exhibited high-quality song, large ornaments, bright or intense colouration, were outbred, with low asymmetry, of a high social rank, and with high-quality territories or nests. I included tests of social rank as long as signalling was recorded in the absence of rivals; this is important because high-rank individuals may suppress the behaviour of subordinates.

#### Body size

This category included studies relating signalling to body length, weight, or some proxy length measurement (e.g. leg length, wing length, pronotum width).

#### Condition

This category included studies examining the effect of: a) diet or food level, b) the relationship between body size and weight, c) direct measurements of body lipid content or plasma metabolite level, and d) environmental conditions that could alter physiological stress in the short-term (oxygen, carbon dioxide, and water acidity in aquatic environments). I used several indirect, morphological measures of condition, though I note that several common measures have been criticised^9,41,42^. I assume that individuals that were relatively heavy for a given body weight, with low lipid stores, or who experienced low food levels, poor-quality diets or stressful environments were in poor condition. I excluded studies examining how signalling behaviour related to physiological markers of stress, as stress responses are typically short-lived and may have a complex relationship with condition (but see^17^).

#### Mated status

This category included studies comparing signalling between virgins and once-mated individuals. I excluded tests related to the number of matings above one, or other forms of mating experience (i.e. the phenotype of previous mating partners).

#### Parasite load

This category included studies measuring the presence or number of external (lice, mites and crustaceans) or internal (acanthocephalans, nematodes, plathyhelminthes, alveolates, fungi, bacteria, and viruses) parasites. I included sexually-transmitted parasites, even in cases where host behavioural changes were hypothesised to be due to parasite manipulation (e.g.^43^). I excluded studies relating behaviour to the presence of endosymbionts in insects, given that the fitness effects on hosts are unpredictable and unknown for most species. Finally, I also excluded studies examining the effect of controlled immune challenges on host behaviour, for example by introducing sterile pellets or inactivated pathogens into the host. This is because any consequences for host condition are indirect in such cases, caused by upregulation of the host immune system, and are typically short-lived (e.g.^44^).

### Effect sizes

I used the correlation coefficient *r* as the effect size in this analysis. Here the effect size represents the *difference* or change in behaviour in relation to individual state. Larger values therefore represent a greater difference in behaviour in relation to state.

I extracted all relevant effect sizes from each paper. This often resulted in multiple effect sizes per paper, either because studies report data relating to multiple experiments, populations or species, or record different behaviours. I obtained effect sizes either: a) directly (if correlations were reported in the text), b) using summary data presented in the study, c) from the results of statistical tests using a range of conversion equations^37,45^, or d) by re-analysing raw data. When studies compared behaviours between multiple categories or treatments (more than at least 3), or across many individuals, I converted the results directly to the correlation coefficient *r*. When studies compared behaviours between two categories or treatments, I first calculated the standardised mean difference, known as Hedges’ *d* ^37,46^, which was then converted to *r*. Hedges’ *d* was calculated either using reported means and standard deviations, or converted from common statistical tests (e.g. t tests, chi-squared, Mann-Whitney U test). I used the online tool WebPlotDigitizer v4 (https://apps.automeris.io/wpd/) to extract raw data from scatter plots, and means and standard deviations from bar plots. Information on methods for effect size calculations are presented in **Table S1**. In cases where effect sizes came from a repeated-measures design (for example using a test statistic from a paired t-test), I assumed a correlation of 0.5 between the two measures of behaviour. In this case, the effect size calculations are identical to those derived from independent-measures tests^37^.

The correlation coefficient can range from +1 to −1. Therefore effect sizes needed to be assigned an arbitrary direction for comparison. Effect sizes were assigned a positive direction when they reflected ability-based signalling, and a negative direction when they reflected needs-based signalling (**Figure 2**). In other words, effect sizes were assigned a positive direction when signalling was higher in individuals that were young, attractive, large, in good condition, and had a low parasite load. Effect sizes were assigned a negative direction when signalling was higher in individuals that were old, unattractive, small, in poor condition, and with a high parasite load. Mated status does not neatly fit into this scheme, so I instead assigned a direction based on the motivation to mate. Here, effect sizes were assigned a positive direction when signalling was higher in virgins (with a high motivation to mate), and negative when signalling was higher in mated individuals. This also reflects the fact that virgins are typically young and in good condition.

**Figure 2.**
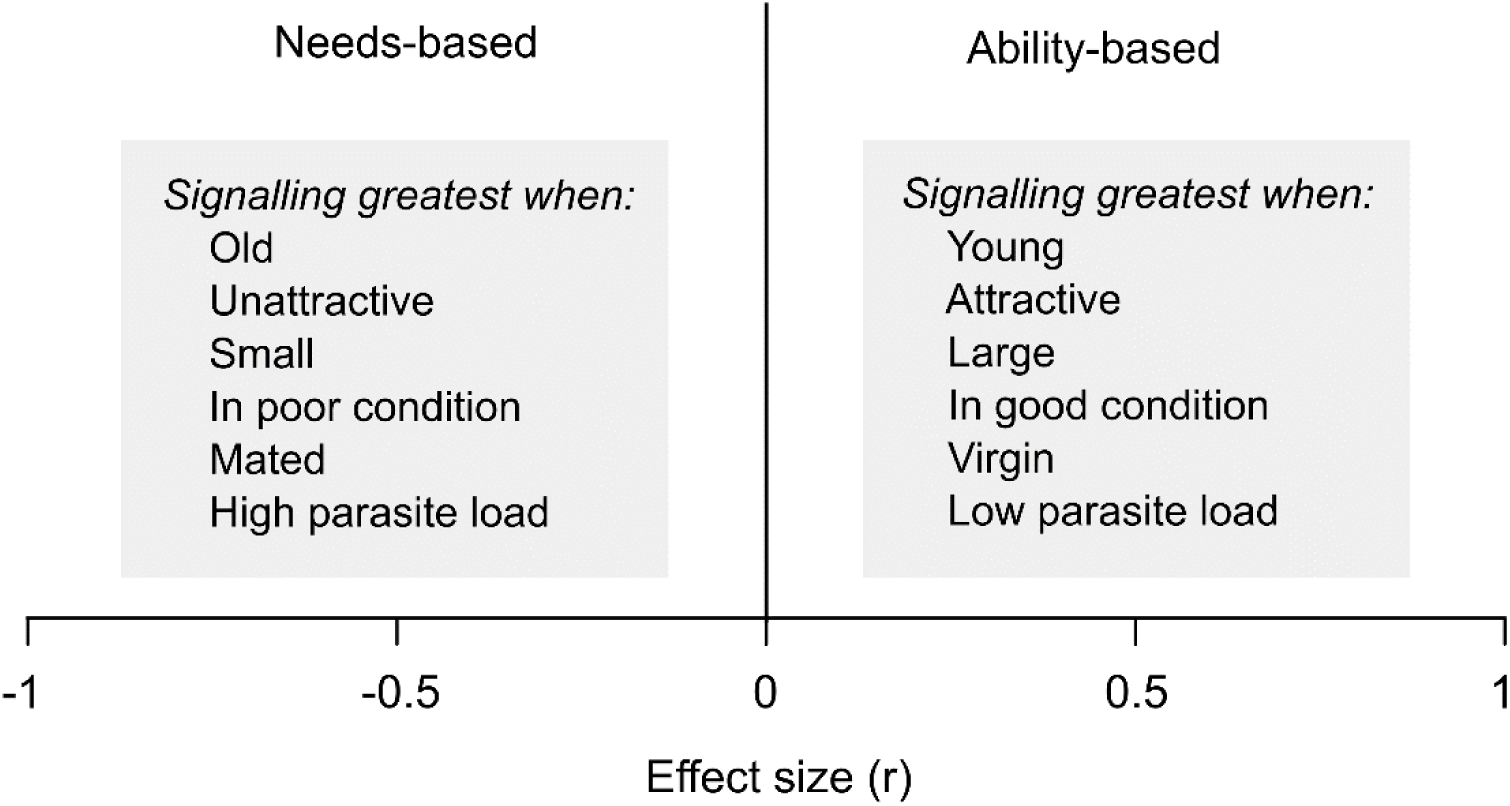
Guide to the coding scheme for effect sizes used in this study. Effect sizes were assigned a positive direction when they reflected ability-based signalling, and a negative direction when they reflected needs-based signalling. Note that the values on the x axis are for illustrative purposes only.

Finally, relevant studies often report non-significant results without describing the direction of the effect. These effect sizes would traditionally be excluded, however this systematically biases the dataset against non-significant results^47^. Therefore, I assigned these effect sizes a value of zero, and ran the analyses with and without including these extra data points. This resulted in two separate datasets: the ‘full’ dataset including all directionless effect sizes, and the reduced dataset with directionless effect sizes excluded (‘no zeroes’ dataset).

### Phylogeny

I incorporated phylogenetic history into all models, in order to account for the potential non-independence of effect sizes from closely-related species. As no single phylogenetic tree was available that included all of the species in my sample, I constructed a supertree using available phylogenetic and taxonomic information using the Open Tree of Life database^48^. Trees were created in R v3.6^49^, using the Rotl^50^ and Ape^51^ packages. For cases where the OTL database resulted in polytomies, I manually searched for published phylogenetic trees for the branches in question. I solved all but one polytomy in this way (the exception is in the *Drosophila* clade). See the online supplementary material for sources. Given the absence of accurate branch length data for these trees, branch lengths were first set to one and then made ultrametric using Grafen’s method^52^. For analyses including subsets of the data I used an appropriately pruned tree. The final ultrametric tree for the full dataset can be seen in the supplementary material (**Figure S1**).

### Moderating factors

For each effect size I additionally recorded information relating to several potential moderators of effect size. I recorded two species-level moderators:

#### Taxonomic group

The class (or equivalent taxonomic level) of the tested species. I predicted that plasticity in signalling behaviour should be highest for long-lived taxonomic groups (mammals, birds, amphibians) which have more than one breeding season over their lifespan. Species that typically only survive one breeding season are predicted to invest maximally into reproduction, and so show reduced plasticity. Short-lived species should also show more negative effect sizes, indicative of terminal investment.

#### Maximum lifespan

I predicted that plasticity in signalling behaviour would be greater for long-lived species. I therefore searched for data on the maximum recorded lifespan of each species in the dataset, in two ways. First, I searched the online databases AnAge^53^, Fishbase (www.fishbase.de), Amphibia Web (www.amphibiaweb.org) and Animal Diversity Web (https://animaldiversity.org/), and the book by Carey & Judge^54^. I found data for 52 species using these reference works. For the remaining species I manually searched online databases for published studies with suitable lifespan estimates. I found data for a further 40 species this way, resulting in data for 92 species in total.

I also recorded a further six effect-size-level moderators:

#### State factor

Whether the degree of plasticity depended on which state factor was tested. If resource-based signalling is important, signalling should be highest for individuals that are young, attractive, large, in good condition, and with a low parasite load. If needs-based signalling is important, signalling should be highest in individuals who are virgin, old, unattractive, in poor condition, and with a high parasite load.

#### State variation

Whether variation in individual state was natural or experimentally-manipulated. I predicted that plasticity would be greater when individuals were experimentally manipulated, as this should increase the ability to statistically detect between-group differences.

#### Signaller sex

Whether signalling was recorded in males or females. I predicted that plasticity in sexual signalling should be stronger for females compared to males, because females generally have a lower reproductive rate and so are more prudent when it comes to allocating resources to reproduction.

#### Study type

Whether signalling behaviour was measured in a laboratory of field setting. I predicted that lab studies should show a stronger relationship between individual state and signalling compared to field studies because: a) field measurements of signalling may be confounded by variable environmental conditions which could also influence the costs and benefits of signalling, b) conditions are typically more benign under laboratory conditions, so that individuals may be able to invest more into reproductive behaviours.

#### Signalling modality

I classed behaviours as either acoustic, olfactory, tactile, visual, or some combination (mixed). I predicted that plasticity should be lowest for olfactory signals, which are energetically cheap, and highest for acoustic and visual signals, which are more energetically expensive.

#### Signalling type

I classified signalling into three main categories. The ‘encounter’ category includes studies which tested signaller behaviour in response to a single encounter with a member of the opposite sex (or some indirect indicator of opposite-sex presence). In this category, behaviour was typically tested once, over a relatively short period, and includes a mix of attraction and courtship signals. The ‘investment’ category includes studies which tested signaller behaviour in the absence of any members of the opposite sex. Here, studies primarily measured medium and long-range attraction signals, and often over a longer time period. Finally, the ‘lek’ category includes species with a lekking mating system, in which members of one sex (typically males) gather and display simultaneously in a small area^55^. In traditional leks, males may signal when members of the opposite sex are close, far away or absent. Studies in this category typically measured lekking behaviour in natural populations, and did not specifically record whether potential mates were present, but this was probably common in most cases. I predicted that plasticity in signalling should be lowest for studies in the ‘encounter’ category, as such displays are likely less costly than the long-range signals seen in the investment and lek categories.

### Statistical analysis

All statistical analyses were performed using R v3.6^49^. Meta-analyses were performed using the package Metafor v2.4^56^. In order to determine the overall mean effect size, I ran a multilevel random-effects model, with study ID, species, phylogeny and observation ID as random factors. Study ID was included as a random factor because I extracted more than one effect size for most studies (mean= 2.28, range= 1-15). Phylogeny was incorporated into the model using a variance-covariance matrix, assuming that traits evolve via Brownian motion. For all analyses, I used Fisher’s *Z* transform of the correlation coefficient (*Zr*), as this has better statistical properties when *r* approaches ±1 ^37^. The associated variance for *Zr* (var *Z*) was calculated as 1/(n - 3) ^57^, with n being the total number of animals used in the test. Model results were then converted back to *r* for presentation. The mean effect size was considered to be significantly different from zero if the 95% confidence intervals did not overlap zero. I first ran this model using the ‘full’ dataset, and then after excluding directionless effect sizes. I used I^2^ as a measure of heterogeneity of effect sizes^58^. I^2^ values of 25, 50 and 75% are considered low, moderate and high, respectively^58^. I calculated I^2^ across all effect sizes, and also partitioned at different levels of the model (study ID, species, phylogeny and observation ID) using the method of Nakagawa & Santos^59^.

I investigated potential moderators of the effect size using the full dataset (i.e. non-directional effect sizes included). To do this I ran meta-regression models, which were identical to the above model except for the inclusion of a single (categorical or continuous) fixed factor. I ran a separate model for each fixed effect. I considered a moderator to significantly influence the mean effect size by examining the *Q*_*M*_ statistic, which performs an omnibus test of all model coefficients. I tested the effect of nine moderators: the eight moderators listed above, plus study year as a test for publication bias (see below). All models were run on the full dataset (k=520), with the exception of lifespan (k= 376). I used the log transformation of lifespan (LnLifespan) because estimates were right-skewed. In order to estimate the average effect size for each level of the categorical moderators, I ran the same meta-regressions as above, but excluding the model intercept (again run separately for each fixed factor).

I searched for signs of two types of publication bias in the full dataset. First, I tested whether the average effect size varied over time, using a meta-regression with study year as a fixed effect (as above). A decrease in effect size over time is a common pattern in novel research fields, and arises when underpowered studies are more likely to be published as a field ages^37^. Second, I searched for funnel plot asymmetry, which can be caused by publication bias against studies with small sample sizes or non-significant results^37^. I tested for funnel plot asymmetry using a trim-and-fill test^60^ and Egger’s regression (regression of Zr against inverse standard error^61^.

## Results

The final (full) dataset consisted of 520 effect sizes from 228 studies and 147 species. The reduced dataset (removing directionless effect sizes) consisted of 399 effect sizes from 184 studies and 122 species. I obtained data from 8 taxonomic groups, though the majority of effect sizes were from insects, fish and birds (**Figure 3a**). The large majority of effect sizes measured male signalling-only 15 effect sizes (from 7 species) considered female signalling (2.9%). Female signalling was recorded for three sex-role reversed fish species (*Syngnathus scovelli, S. typhle* & *Culaea inconstans*), one partially role-reversed bushcricket (*Requena verticalis*), one bird species with biparental care (*Taeniopygia guttata*), one amphibian with both male and female calls (*Eleutherodactylus cystignathoides*), and a bug with female-produced attractant pheromones (*Trigonotylus caelestialium*). Of the six state factors examined, the majority (70%) of effect sizes related to body size and condition (**Figure 3a**). Overall, sexual signalling was positively related to individual state, so that signalling was significantly higher in individuals with greater resources (Mean r= 0.127, 95% CI= 0.074-0.178, k= 520; **Figure 3b**). However, the mean estimate was small^62^. Removing the 121 directionless effect sizes led to a slight increase in the mean effect size (Mean r= 0.168, 95% CI= 0.102-0.231, k= 399). The full dataset showed high total heterogeneity (Total I^2^= 90.68%), and examination of the funnel plot shows large variation in both positive and negative effect sizes (**Figure 3b**). A low-moderate proportion of total heterogeneity was attributable to between-study differences (19.53%) and between-species differences (26.67%). Phylogenetic history explained a negligible amount of heterogeneity in effect sizes (0%).

**Figure 3.**
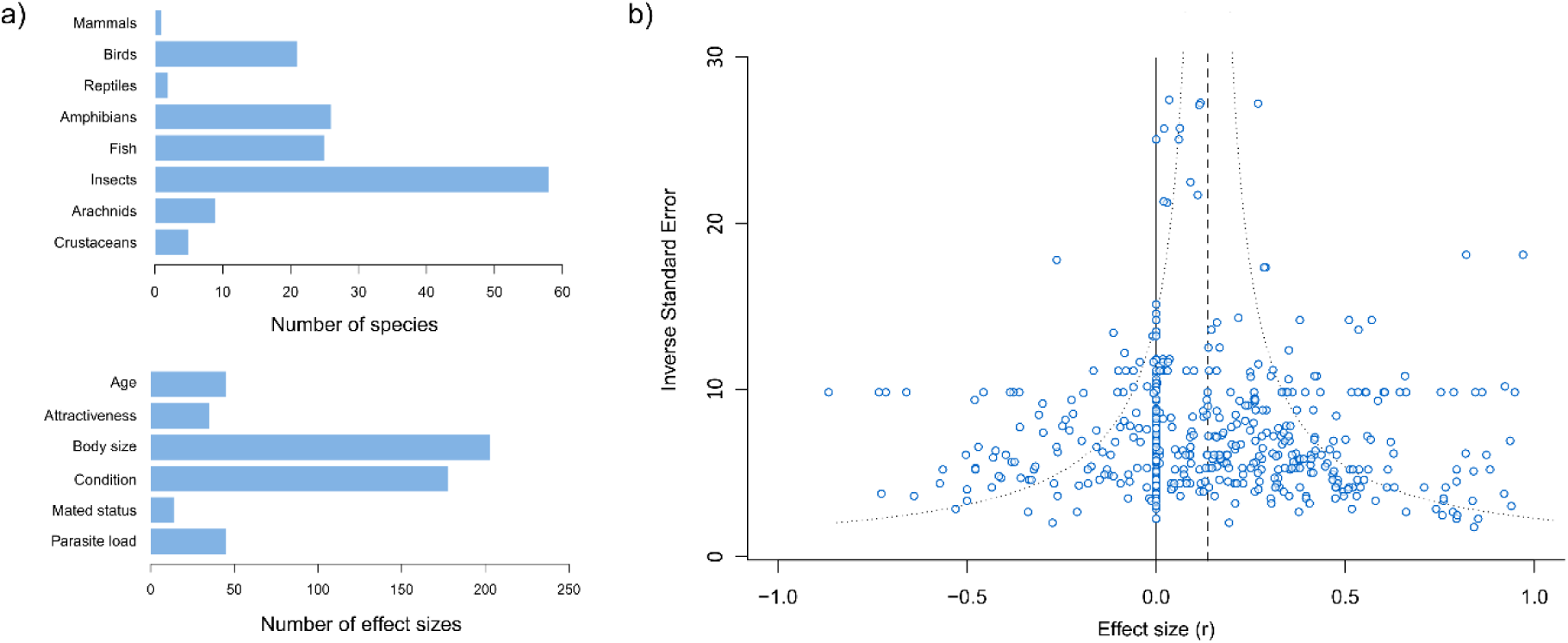
The final dataset consists of 520 effect sizes from 228 studies and 147 species. **a)** Histograms showing the number of species in each taxonomic group category (top panel) and the number of effect sizes in each state factor category (bottom panel). **b)** Funnel plot showing the relationship between effect size (the correlation coefficient r) and the inverse standard error (a measure of study precision-larger values represent studies with larger sample sizes) for the full dataset (k= 520, directionless effect sizes included). The dashed line shows the overall mean effect size estimate from a multilevel random-effects meta-analysis model (see text for details). The dotted line illustrates the typical expected ‘funnel’ shape, with large sample-size studies resulting in effect size estimates closer to the mean.

The strength and direction of behavioural plasticity was significantly influenced by which aspect of individual state was measured (**Table 2**). The mean estimates for all state factors were positive with the exception of age (**Figure 4, Table S2**). However, only age and condition had mean estimates that differed significantly from zero: signalling was greatest when individuals were old and in good condition (**Figure 4, Table S2**). There was also a marginally-significant effect of body size, with signalling greatest for large individuals (**Figure 4, Table S2**). The strength of plasticity was also greater when state was experimentally manipulated, compared to when state varied naturally (**Table 2; Table S2**). The strength and direction of behavioural plasticity was not significantly influenced by taxonomic group, sex, the type of signalling, lifespan, or any of the other moderator variables tested (**Table 2; Table S2**).

**Table 2.** Results from meta-regressions testing the nine moderator variables. Each factor was tested using a multilevel mixed-effects meta-analysis model (meta-regression), with study ID, species, phylogeny and observation ID as random factors, and a single continuous or categorical fixed factor. The *Q*_*M*_ statistic tests whether the moderator variable significantly influences the mean effect size. All models were run on the full dataset (k=520), with the exception of lifespan (k= 376). Significant moderators are highlighted in bold.

| *Factor* | *Q<sub>M</sub>* | *P* |
| --- | --- | --- |
| Taxonomic class | 8.89 | 0.26 |
| Study type | 0.47 | 0.49 |
| Signaller sex | 0.22 | 0.64 |
| **State factor** | **32.62** | **<0.001** |
| **State variation** | **6.44** | **0.01** |
| Signalling modality | 4.12 | 0.39 |
| Signalling type | 2.7 | 0.26 |
| LN(Lifespan) | 0.07 | 0.79 |
| Study year | 2.12 | 0.15 |

**Figure 4.**
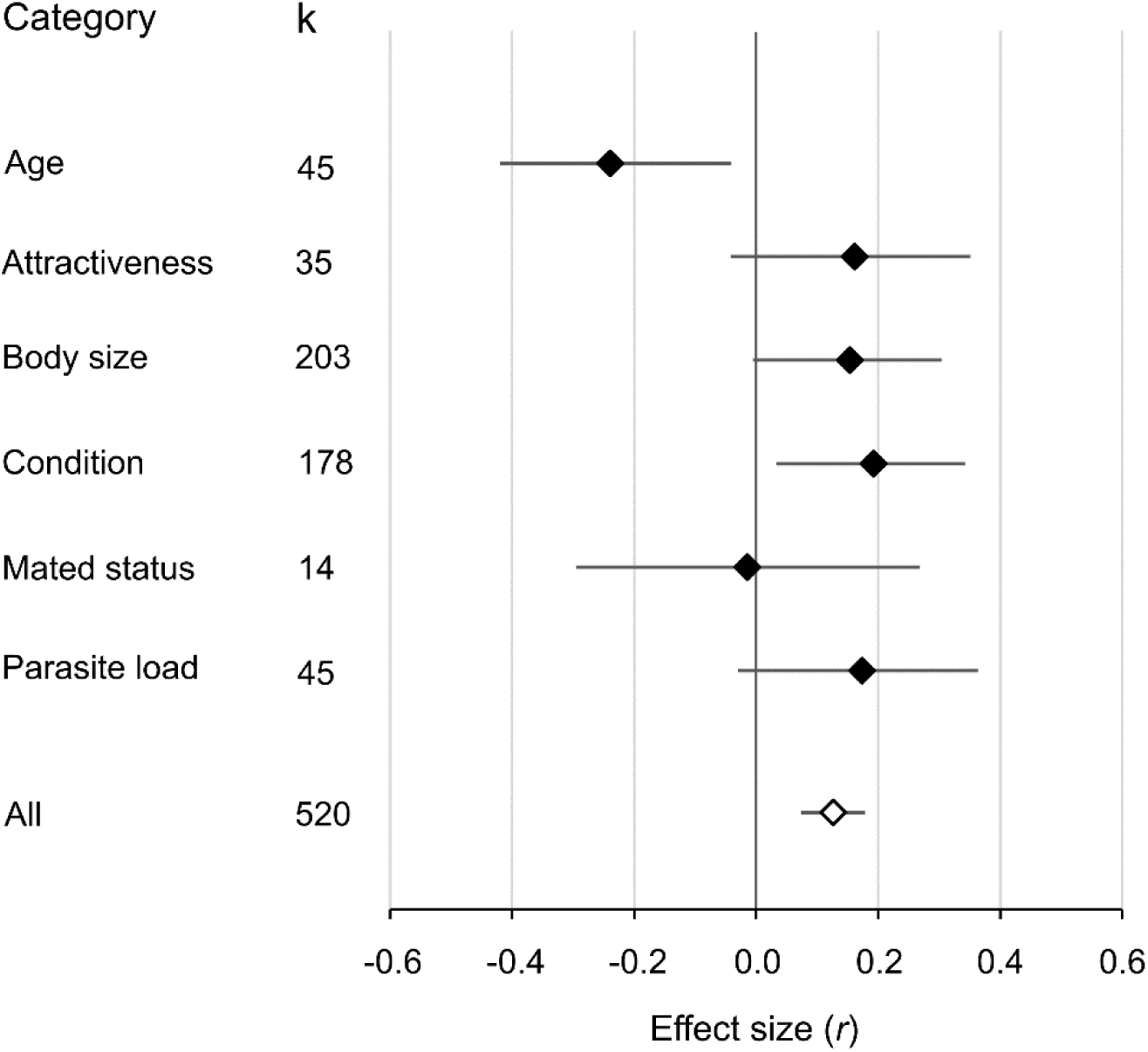
Forest plot showing the mean effect size (the correlation coefficient *r*) estimate for each state factor (black symbols). Estimates were obtained from a meta-regression model with the intercept removed (see text for details). The mean effect size estimate for all effect sizes is shown for comparison (open symbol). In all cases, diamonds represent the mean estimate and the bars represent the 95% confidence intervals. K= the number of effect sizes for each category.

The mean effect size was not significantly related to study year (**Table 2, Figure S2**). There was no significant relationship between effect size and inverse standard error (*F*_1,518_= 1.98, *P*= 0.16; **Figure S3**). A trim-and-fill test detected significant funnel plot asymmetry. However, all 82 missing effect sizes were detected to the right of the mean estimate, so they are unlikely to be due to publication bias against non-significant or negative results. Adding these ‘missing’ effect sizes to the dataset resulted in a more positive overall estimate (Mean r= 0.217, 95% CI= 0.189-0.245; **Figure S4**).

## Discussion

Across 228 studies and 147 species, animal sexual signalling was significantly influenced by individual state, with individuals (who in almost all cases were male) generally signalling more when in good condition. Overall, this suggests that sexual signalling behaviour is weakly honest, and dependent on current available resources (ability-based). There was little evidence for consistent needs-based investment in sexual signalling, except when considering old virgins, who exhibited greater signalling when compared to young virgins. Further, when considering the six state factors separately, only condition and age had mean estimates that were significantly different from zero. However, there was a strong trend for individuals to signal more when attractive, large, and with a low parasite load. Finally, sexual signalling behaviour was not significantly influenced by mated status, though the sample size for this category was small.

In this dataset, effect sizes were coded so that negative values represented needs-based signalling (or terminal investment). Examination of the funnel plot show many examples of such needs-based signalling in the form of negative effect sizes, including several of large magnitude. Needs-based signalling was not more common in short-lived species, either when comparing taxonomic groups or with maximum lifespan data. There was also little evidence that parasites manipulate their host’s signalling behaviour in order to increase transmission^63,64^. However, consistent needs-based signalling was seen in those studies examining individual age, with signalling being significantly greater in old compared to young individuals, as predicted by the terminal investment hypothesis. However, it is important to note that I only included studies testing age-dependent signalling of virgins in this category, in order to control for the confounding effect of mated status on signalling behaviour. Further, all effect sizes in this category considered male signalling. This means that this result will only be ecologically realistic for populations in which a large proportion of sexually-mature males remain virgins, such as those in which male mating success is highly skewed. Nevertheless, it does fit with the idea that terminal investment is only favoured when residual reproductive value is reduced to a large extent^25^, such as when trying to avoid total mating failure. Similarly, virgin female pheromone signalling also increases with age across 44 moth species^19^. It would be informative to compare these results to cases where sexual signalling is measured in mated individuals. I predict that in such studies signalling will instead be highest for young individuals, in line with the results for the majority of this dataset.

While significantly positive, the overall effect size across studies was small^62^, even after removing a large number of directionless effect sizes. This can be attributed partly to the large amount of heterogeneity observed across studies. Some of this variation was explained by experimental design: plasticity was greater in studies where individual state was experimentally manipulated, compared to those examining natural variation in state. This is probably because experimental manipulations tend to reduce within-group variation, and so increase the statistical power to detect differences between groups. This result, combined with the ability to confirm causal relationships^9^, means that experimental studies will be more useful in future, and should be encouraged. Nevertheless, the majority of effect size heterogeneity remains unexplained. Notably, a negligible amount of heterogeneity was attributed to phylogenetic history, suggesting that behavioural plasticity is not evolutionarily constrained. In contrast, almost half (44%) of this variation is at the observation level. This variation could arise for example due to within-study differences in signalling behaviours, the conditions experienced by signallers prior to tests, or the error associated with imperfect measures of individual state^41,42^. Additionally, two potentially important moderators which may explain variation in the magnitude of plasticity, and which could not be incorporating into this analysis, are the relative energetic or fitness cost of signal expression, and the range of individual states measured. Plasticity should be greatest for signals that are costly to produce, and for studies examining a wide range of individual states. However, these factors will likely prove very difficult to quantify and compare in a standardized way across species.

This analysis is limited in three important ways. First, I only tested for linear relationships between state and sexual signalling, because of the difficulty in identifying ‘intermediate’ categories in a standardised way. However, for some aspects of state, quadratic effects may be important. For example, signalling is often predicted to be highest at intermediate ages^19^, possibly because of a balance between resource level and residual reproductive value. Second, I only examined quantitative variation in specific display behaviours. Consequently, there are a range of other forms of behavioural plasticity that are not included in this dataset. For example: small, unattractive or poor-condition males may adopt discrete alternative mating tactics, such as sneaking or female mimicry, instead of signalling^65^. Males may also adapt their displays in more subtle ways. For example, male Endler’s guppies strategically present females with their most attractive side during courtship^66^. Third, the dataset is extremely male-biased: less than 3% of effect sizes examined female signalling. Though in general males are more likely to search for and signal to mates, female sexual signals are not excessively rare, and are very widespread in some clades (e.g. Lepidoptera, birds). Indeed, the meta-analysis by Umbers *et al*.^19^ examining age-related variation in virgin female pheromone signalling in moths found 38 relevant papers, none of which were picked up by my search criteria, probably due to the absence of the phrase ‘pheromone’ in my searches. This highlights the fact that the sample analysed here is by no means exhaustive. Nevertheless, more studies in females are needed in order to make an informative comparison between the sexes. This is important because some of the factors examined here, such as mated status, are predicted to more strongly influence the expression of female reproductive behaviour when compared to males.

Overall, individuals generally signal more when they have the resources available to do so. However, determining the causal relationships between individual state and sexual signalling may be difficult, for two reasons. First, explanations based on the costs or benefits of signalling sometimes predict plasticity in the same direction, and most (correlational or experimental) studies cannot exclude one or the other. For example, good-condition individuals could signal more because they can afford the costs of signalling, or because a given signalling bout is more likely to be successful. Future studies will need to design experimental manipulations that can control for one of these explanations. Second, many of the state factors examined here may be correlated with each other. For example, age is often correlation with body size and condition, and attractiveness may be correlated with body size, condition or parasite load. While some studies control for these correlations, others do not (and this is especially true for correlational studies). Both of these problems emphasise the need for more careful experimental manipulations of individual state.

## Acknowledgements

I would like to thank Tom Price and Zen Lewis for helpful comments on an earlier version of the manuscript.

## Author contributions

LRD conceived the study, carried out the searches, collected the data, performed the analyses, and wrote the manuscript.

## Supplementary background

The six state factors examined here have been theorised to influence sexual signalling in a variety of ways.

### Age

The relationship between age and sexual signalling is predicted to be complex, as the fitness payoffs associated with signalling could vary with age for several reasons. The resources available to an individual may increase with age for many species^1^. Further, age may also be correlated with other important determinants of mating success, such as body size (e.g. in species with indeterminate growth^2^) and dominance^3^. However, old individuals are often in poorer condition that young individuals, due to age-related declines in physiology^4,5^. Therefore, condition and competitiveness may be highest in young or old individuals, or even somewhere in between, across species. Alternatively, the terminal investment hypothesis predicts that old individuals should invest more into sexual signalling, because they face fewer future mating opportunities than young individuals^6,7^. The opportunity for future reproduction is also referred to as ‘residual reproductive value’, with younger individuals having a higher value than older individuals, and so benefit more by conserving resources^8^. Importantly, age is often confounded by mated status, especially for wild individuals, and this may influence the motivation to signal independently of age.

### Mated status

Mated status (whether individuals are virgin or mated) strongly influences the motivation to mate: virgins should invest heavily in signalling in order to achieve their first mating^9^, as failing to do so results in a fitness of zero. Because fitness typically increases with each mating, each subsequent mating is less beneficial for mated individuals. This means mated individuals can be more strategic with their signalling compared to virgins.

### Attractiveness

Attractive individuals are those that possess phenotypes that are preferred by members of the opposite sex, resulting in above-average mating success. Attractive individuals may be of higher ‘quality’, in terms of the direct or indirect benefits they can provide mates^10,11,12^, or better able to exploit the sensory biases of choosers^13^. The relationship between attractiveness and signalling effort may be mediated by several processes. First, attractiveness may be correlated with condition if mate choice is based on condition-dependent traits^14^, so that attractive individuals can afford to signal more (but see ^15^). Second, the payoffs of signalling may often be higher for attractive individuals, in the sense that a given signalling bout is more likely to be successful (differential benefits hypothesis^16,17^), again suggesting that attractive individuals should signal more. In such cases it may pay unattractive males to invest in alternative mating tactics, such as sneaking behaviour, rather than invest heavily in display behaviour that has a low chance of success^18^. Alternatively, in some cases attractiveness may influence the cost of signalling. For example, large (attractive) male guppies have a greater predation risk, and so may signal less in some contexts^16^. Finally, the relationship between attractiveness and signalling may also depend on how attractiveness is assessed by the other sex. If attractiveness is assessed partly by traits other than behavioural displays, then attractive individuals may be able to afford to display less, while investing in morphology or colouration for example^16^. However, when attractiveness is determined based on display behaviour, then all individuals should display as much as they can afford, irrespective of their attractiveness.

### Body size

Body size may influence signalling behaviour in multiple ways. Larger individuals may be in better condition, and so be better able to pay the energetic costs of signalling. Larger individuals are often more attractive, and better able to displace rivals, and so may also benefit more from signalling than smaller individuals^16,17^. Finally, body size also has the potential to influence the benefits of signalling, by influencing signal production or detection. For example, larger individuals may produce louder or deeper calls because of differences in the size of sound-producing organs^19,20^. However, for this reason signalling could be costlier for large individuals, for example if louder calls lead to an increased predation risk^21^.

### Condition

Condition is generally ill-defined^22^. However, when considered in the context of signalling behaviour, condition typically refers to the amount of resources an individual is able to allocate to reproduction^23^. Therefore, poor-condition individuals are expected to have fewer energetic reserves available to invest in sexual signalling compared to high-condition individuals. Additionally, poor-condition individuals may also have fewer future mating opportunities, because of a reduction in lifespan or attractiveness, and so could benefit more from signalling^7^. Importantly, in contrast to the other factors considered here, condition and sexual signalling are directly linked, due to the fact that the expression of costly signals will unavoidably reduce the available resource pool. Therefore, signalling and condition are causally linked, so that individuals could be in poor condition *because* they signal more. Experimental studies are therefore needed in order to determine the direction of causality^24^.

### Parasite load

Parasites may reduce the amount of resources available, either because of direct parasite effects or costly upregulation of the host immune system^7^, and so parasite effects are often seen as a special case of condition-dependence^25^. Therefore parasitism is generally expected to reduce host signalling behaviour. Parasites may also influence attractiveness, especially if they interfere with the production of ornaments, for example by damaging plumage^26^ or disrupting host hormones (the ‘immunocompetence handicap’ hypothesis^27^); and this could also influence signalling behaviour (see above). However, parasites could also manipulate hosts to increase signalling^28,29^, in order in order to facilitate transmission through predation (if predators are an intermediate host) or mating (if parasites are transmitted sexually).

## Online database searches

I performed literature searches using the online databases Web of Science & Scopus on the 13/08/2019. I used the following search terms relating to mating behaviour:

- (mate OR mating) AND (choice OR preference* OR choosiness OR rejection)
- courtship OR courting OR “sexual signalling”
- mate AND (sampl* OR search*)
- ”species recognition” OR “mate recognition” OR “reproductive isolation” OR (conspecific* AND discriminat*) OR ((mate OR mating) AND (hybridisation OR reinforcement)) OR (mating AND (heterospecific* OR interspecific*))

I used the following search terms relating to individual state:

- age OR “mated status” OR “mating status” OR “mating history” OR “number of matings” OR virgin* OR parasite* OR disease OR diet OR hunger OR food OR stress OR condition OR size OR weight OR quality OR attractiveness OR resource* OR “territory quality” OR “reproductive cycle” OR “social rank” OR inbreeding OR personality OR boldness OR exploration OR “behavioural syndrome*”
- NOT human

Search results were imported into Endnote (Clarivate Analytics), and duplicates removed. All searches combined resulted in 32320 results. I then screened the titles using Endnote to remove obviously irrelevant studies (e.g. studies on humans, other subjects, review articles; Figure 1). This resulted in 7158 studies. Next I imported all of the abstracts into Rayyan for further screening. This resulted in 1230 promising studies, which were downloaded and read in their entirety.

**Table S1.** Methods for re-analysis of study data

| Study | Source | Test |
| --- | --- | --- |
| Adamo et al., 2014 | Text | Chi-squared |
| Ahtiainen et al., 2006 | Text | Hedges' d |
| Amorim et al., 2013 | Figure 4 | Spearman's correlation |
| An & Waldman, 2016 | Figure 2 | Hedges' d |
| Anderson & Jones, 2019 | Figure 5 | Correlation |
| Andrade & Mason, 2000 | Text | Chi-squared |
| Anichini et al., 2018 | Figure 3 | Hedges' d |
| Atagan & Forstmeier, 2012 | Table 2 | Hedges' d |
| Bakker & Milinski, 1991 | Figure 3 | Hedges' d |
| Becker et al., 2012 | Figure 1 | Hedges' d |
| Bertram et al., 2009 | Figure 1 | Hedges' d |
| Bertram et al., 2009 | Table 1 | Chi-squared |
| Birkhead et al., 1998 | Table 1 | Hedges' d |
| Buchanan et al., 2003 | Figure 4 | Hedges' d |
| Candolin & Salesto, 2009 | Figure 1 | Hedges' d |
| Chemnitz et al., 2015 | Figure 1 | Hedges' d |
| Clark et al., 1997 | Figure 2 & 3 | Hedges' d |
| Clayton, 1990 | Figure 5 | Mann-Whitney |
| Cobb et al., 1987 | Table 1 | Hedges' d |
| Cordes et al., 2015 | Figure 2 | Hedges' d |
| Devigili et al., 2013 | Table 1 | Hedges' d |
| Drayton et al., 2010 | Table 1 | Hedges' d |
| Droney, 1998 | Figure 3 | Correlation |
| Eastwood & Burnet, 1977 | Table 1 | Hedges' d |
| Engqvist & Sauer, 2003 | Table 1 | Hedges' d |
| Greenspan et al., 2016 | Table 1 | Hedges' d |
| Gumulka & Rozenboim, 2017 | Figure 1 | Hedges' d |
| Hankison & Ptacek, 2007 | Text | Hedges' d |
| Hankison & Ptacek, 2007 | Figure 1 | Hedges' d |
| Head et al., 2017 | Figure 3 | Hedges' d |
| Head et al., 2017 | Figure 4 | Spearman's correlation |
| Hegde & Krishna, 1997 | Table 2 | Hedges' d |
| Henderson et al., 2018 | Raw data | Mann-Whitney |
| Hermann et al., 2015 | Figure 4 | Wilcoxon signed-rank |
| Hoefler et al., 2010 | Figure 1 | Chi-squared |
| Hoefler et al., 2009 | Figure 1 | Hedges' d |
| Holzer et al., 2003 | Text | Hedges' d |
| Honarmand et al., 2015 | Table 3 & 4 | Hedges' d |
| Houde & Torio, 1992 | Figure 2 | Hedges' d |
| Houslay et al., 2017 | Raw data | Spearman's correlation |
| Jacot et al., 2004 | Figure 2 | Hedges' d |
| Jacot et al., 2008 | Text | Hedges' d |
| Jennions, 1998 | Table 3 | Hedges' d |
| Jha & Kumar, 2016 | Figure 1 | Hedges' d |
| Kaspi et al., 2000 | Figure 3 | Hedges' d |
| Kim & Choe, 2003 | Table 1 | Chi-squared |
| Kim et al., 2008 | Figure 4 | Spearman's correlation |
| King et al., 2005 | Table 1 & 4 | Chi-squared |
| Kitsunezuka et al., 2013 | Figure 1 | Chi-squared |
| Kodric-Brown, 1989 | Table 1 | Hedges' d |
| Kolluru et al., 2014 | Figure 5 | Correlation |
| Kotiaho, 2000 | Figure 2 | Hedges' d |
| Kotiaho, 2002 | Figure 2 | Hedges' d |
| Kotiaho et al., 2001 | Figure 2 | Hedges' d |
| Krishna & Hegde, 2003 | Table 5 & 6 | Hedges' d |
| Kuczynski et al., 2015 | Figure 1 | Correlation |
| Kuczynski et al., 2016 | Figure 1 | Correlation |
| Lehtonen et al., 2016 | Figure 3 | Hedges' d |
| Lehtonen et al., 2018 | Figure 5 | Hedges' d |
| Lomborg & Toft, 2009 | Figure 2 | Spearman's correlation |
| Lubanga et al., 2016 | Figure 2 | Spearman's correlation |
| Magellan et al., 2005 | Figure 2 | Hedges' d |
| Magoolagan et al., 2018 | Figure 2 | Spearman's correlation |
| Manica et al., 2014 | Figure 1 | Spearman's correlation |
| Mappes et al., 1996 | Figure 1 | Hedges' d |
| Mariette et al., 2006 | Figure 1 | Hedges' d |
| Mateos & Carranza, 1999 | Text | Chi-squared |
| Matsuo, 2018 | Figure 2 & 4 | Hedges' d |
| McAuley & Bertram, 2016 | Figure 4 | Spearman's correlation |
| McNeil et al., 2016 | Figure 3 | Hedges' d |
| Memmott & Briffa, 2015 | Figure 2 & 4 | Spearman's correlation |
| Milazzo et al., 2016 | Text | Hedges' d |
| Murphy, 1999 | Text | Hedges' d |
| Nickley et al., 2016 | Figure 1, 2 & 3 | Hedges' d |
| O'Hanlon et al., 2018 | Figure 2 | Spearman's correlation |
| Olsson et al., 2009 | Figure 1c | Hedges' d |
| Ortiz-Santaliestra et al., 2009 | Table 1 | Chi-squared |
| Papadopoulos et al., 1998 | Figure 3 | Hedges' d |
| Pariser et al., 2010 | Figure 2 | Hedges' d |
| Pelabon et al., 2005 | Text | Hedges' d |
| Prathibha et al., 2011 | Table 2 | Hedges' d |
| Pryke & Andersson, 2005 | Table 3 | Hedges' d |
| Ptacek & Travis, 1996 | Figure 2, 3 & 4 | Spearman's correlation |
| Rahman et al., 2014 | Figure 1 | Hedges' d |
| Rahman et al., 2013 | Table 4 | Hedges' d |
| Reifer et al., 2018 | Text | Hedges' d |
| Rezaei et al., 2015 | Table 2 | Hedges' d |
| Richardson & Smiseth, 2019 | Figure 3 | Hedges' d |
| Ritchie et al., 1998 | Figure 3b | Mann-Whitney |
| Ritschard & Brumm, 2012 | Figure 3 | Hedges' d |
| Scheuber et al., 2003 | Figure 3 | Hedges' d |
| Shakeel et al., 2010 | Figure 1 | Chi-squared |
| Shelly & Kennelly, 2003 | Table 2 | Hedges' d |
| Sikkel, 1995 | Table 2 | Chi-squared |
| Simmons, 1994 | Table 2 | Hedges' d |
| Sisodia & Singh, 2004 | Table 4 | Hedges' d |
| Snekser et al., 2009 | Figure 1 | Hedges' d |
| Sundin et al., 2015 | Table 3 | Hedges' d |
| Suzaki et al., 2015 | Figure 1 | Correlation |
| Takeshita et al., 2018 | Text | Hedges' d |
| Wacker et al., 2012 | Figure 2d | Chi-squared |
| Whattam & Bertram, 2011 | Table 1 | Hedges' d |
| Win et al., 2013 | Table 1 | Hedges' d |
| Woodhead, 1986 | Table 2 | Hedges' d |
| Yamada & Soma, 2016 | Figure 2a | Hedges' d |
| Yamane & Yasuda, 2014 | Figure 1 | Hedges' d |

### Phylogeny

For the relationships among the Austruca species I used Shih *et al*.^30^. For the relationships among the Lycosidae I used Piacentini & Ramirez^31^. For the relationships among Schizocosa I used Stratton^32^. For the relationships among the Tettigoniidae I used Mugleston *et al*.^33^. For the relationships among the Gryllidae I used Chintauan-Marquier *et al*.^34^. For the relationship among Gryllus species I used Huang *et al*.^35^. For the relationship between beetle orders I used Zhang *et al*.^36^. For the relationships among Onthophagus species I used Emlen *et al*.^37^. For the position of *Drosophila malerkotliana* I used Van der Linde *et al*.^38^.

---

**Figure S1.**
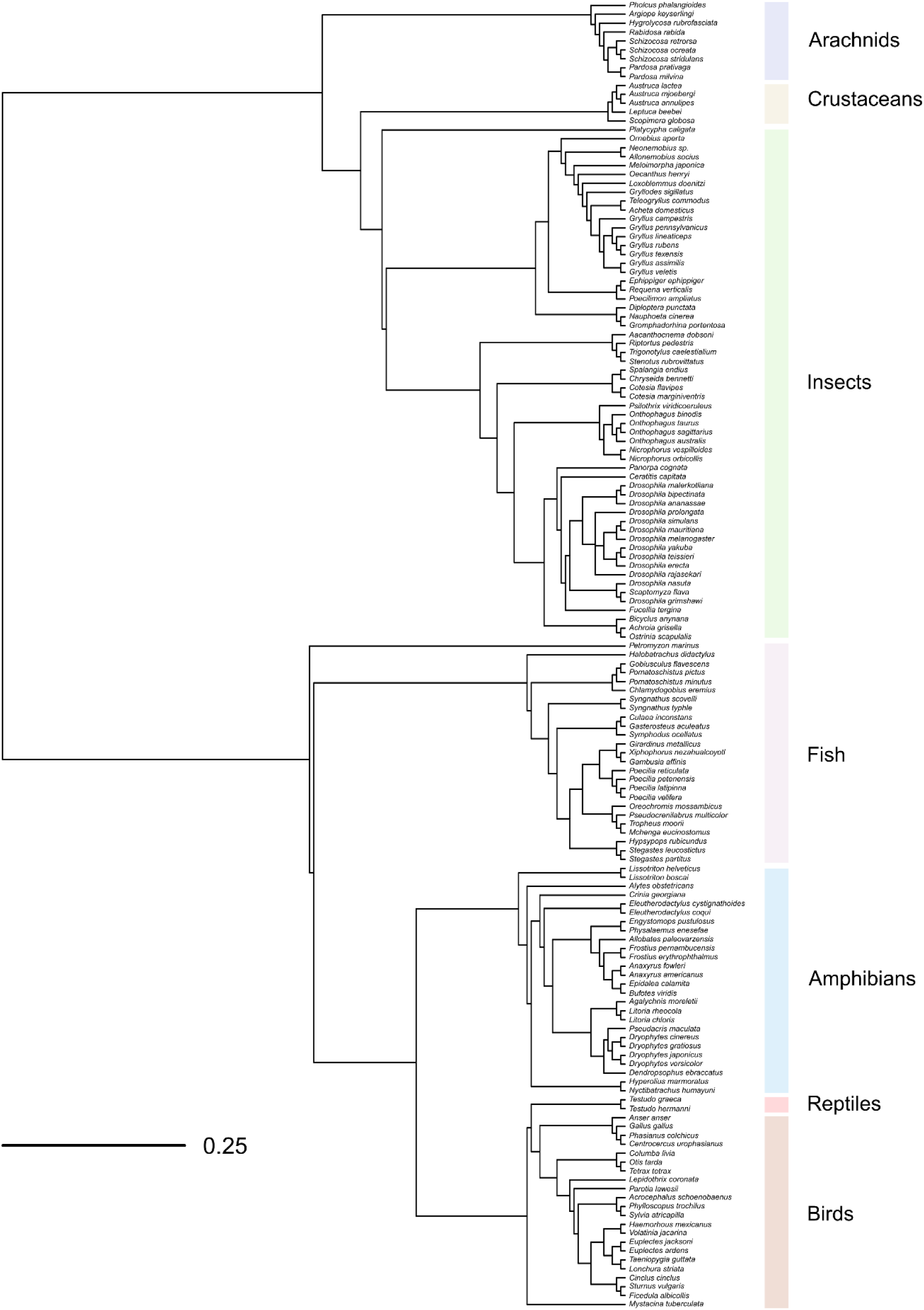
The phylogenetic tree used in the analysis (N= 147 species). The taxonomic classes used during meta-regressions are highlighted on the right-hand side of the tree.

## Supplementary results

**Table S2.** Mean effect size estimates (the correlation coefficient r), plus sample sizes, for each category for the seven categorical moderator variables. Estimates were obtained using a minus-intercept meta-regression, performed separately for each moderator (see text for details).

| Moderator | Category | N Effect sizes | N Studies | N Species | Mean $r$ | 95% CI Lower | 95% CI Upper |
| --- | --- | --- | --- | --- | --- | --- | --- |
| Taxonomic class | Mammals | 2 | 1 | 1 | -0.38 | -1.12 | 0.36 |
|  | Birds | 73 | 39 | 21 | 0.22 | 0.07 | 0.37 |
|  | Reptiles | 4 | 2 | 2 | 0.00 | -0.50 | 0.50 |
|  | Amphibians | 70 | 32 | 26 | 0.03 | -0.10 | 0.16 |
|  | Fish | 122 | 48 | 25 | 0.18 | 0.06 | 0.30 |
|  | Insects | 189 | 79 | 58 | 0.09 | 0.01 | 0.18 |
|  | Arachnids | 42 | 18 | 9 | 0.23 | 0.04 | 0.42 |
|  | Crustaceans | 18 | 9 | 5 | 0.22 | -0.04 | 0.48 |
| Study type | Field | 147 | 70 | 55 | 0.15 | 0.06 | 0.24 |
|  | Lab | 373 | 160 | 98 | 0.12 | 0.05 | 0.18 |
| Signaller sex | Female | 15 | 8 | 7 | 0.07 | -0.16 | 0.31 |
|  | Male | 505 | 223 | 144 | 0.13 | 0.08 | 0.18 |
| State factor | Age | 45 | 19 | 21 | -0.24 | -0.45 | -0.04 |
|  | Attractiveness | 35 | 25 | 20 | 0.16 | -0.04 | 0.37 |
|  | Body size | 203 | 97 | 83 | 0.15 | -0.01 | 0.31 |
|  | Condition | 178 | 104 | 62 | 0.19 | 0.03 | 0.36 |
|  | Mated status | 14 | 7 | 7 | -0.01 | -0.30 | 0.27 |
|  | Parasite load | 45 | 18 | 17 | 0.18 | -0.03 | 0.38 |
| State variation | Manipulated | 146 | 84 | 48 | 0.21 | 0.13 | 0.30 |
|  | Natural | 374 | 156 | 123 | 0.09 | 0.04 | 0.15 |
| Signalling modality | Acoustic | 213 | 110 | 73 | 0.12 | 0.04 | 0.19 |
|  | Mixed | 53 | 25 | 28 | 0.07 | -0.06 | 0.20 |
|  | Olfactory | 29 | 12 | 7 | -0.02 | -0.24 | 0.20 |
|  | Tactile | 28 | 14 | 15 | 0.13 | -0.02 | 0.28 |
|  | Visual | 197 | 88 | 56 | 0.18 | 0.10 | 0.26 |
| Signalling type | Encounter | 296 | 133 | 89 | 0.15 | 0.08 | 0.22 |
|  | Investment | 196 | 84 | 60 | 0.08 | 0.00 | 0.16 |
|  | Lek | 28 | 14 | 7 | 0.24 | 0.01 | 0.48 |

**Figure S2.**
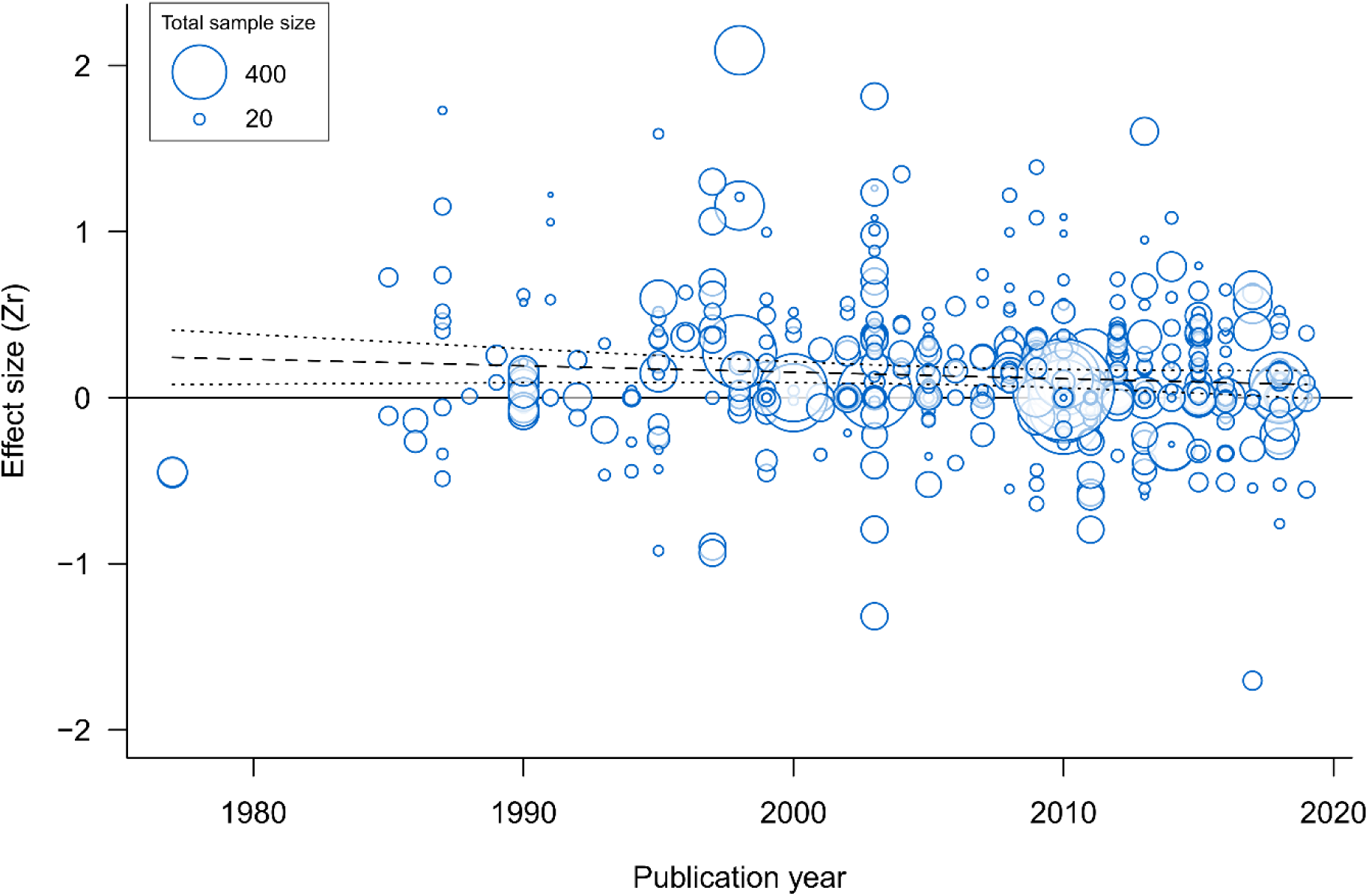
Relationship between effect size (Z-transformed correlation coefficient, *Zr*) and study year. Each bubble represents an effect size, and bubble size is scaled to effect size precision (inverse standard error; larger bubbles reflect larger sample sizes) for the full dataset (k= 520). The dashed line shows the predicted line from a meta-regression including study year as a covariate (see text for details). Dotted lines show the 95% confidence intervals for the predicted line.

**Figure S3.**
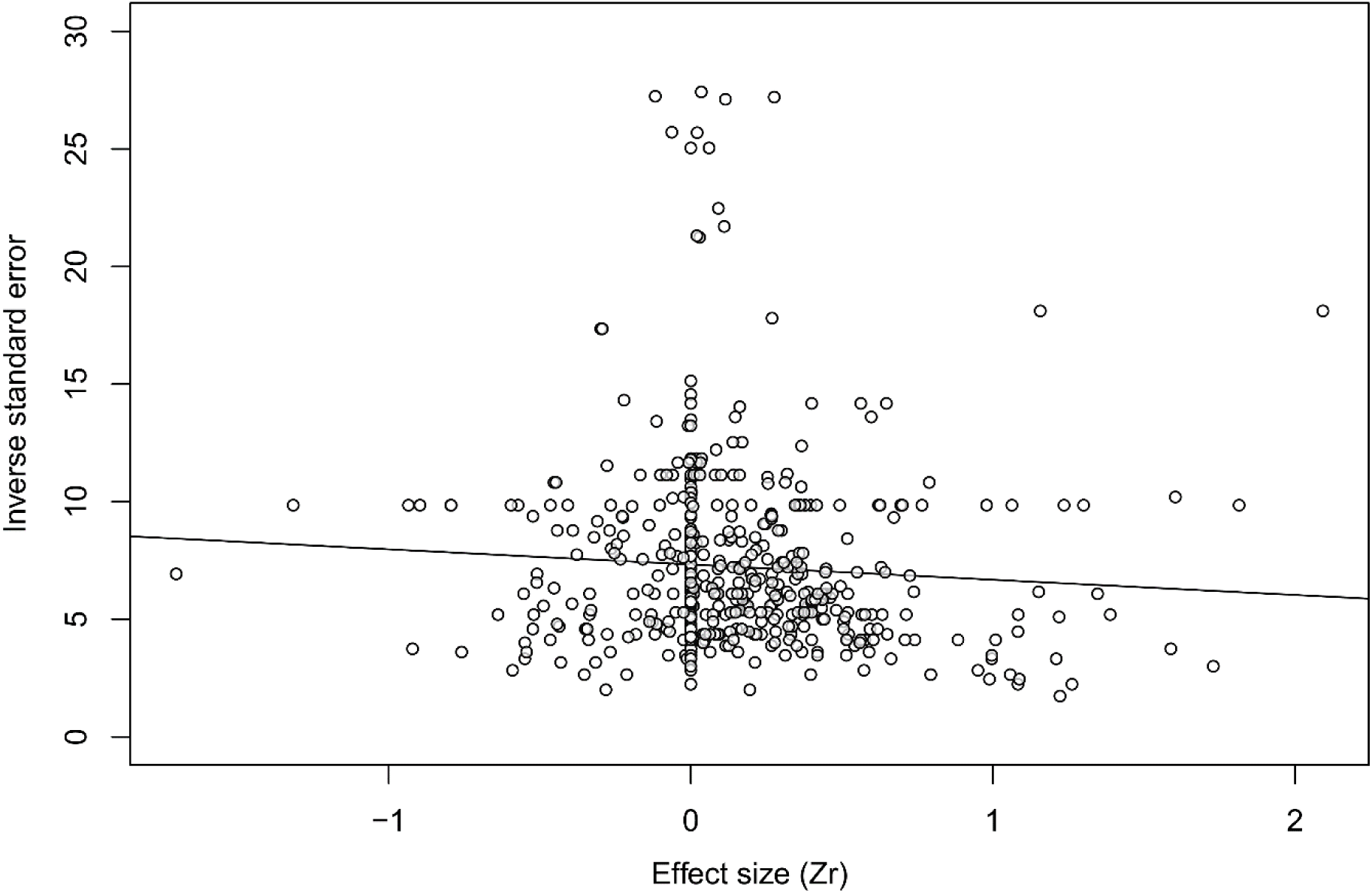
Scatterplot showing the relationship between effect size (Z-transformed correlation coefficient, *Zr*) and precision (inverse standard error; larger bubbles reflect larger sample sizes) for the full dataset (k= 520). The line shows the prediction from a linear regression (Egger’s regression).

**Figure S4.**
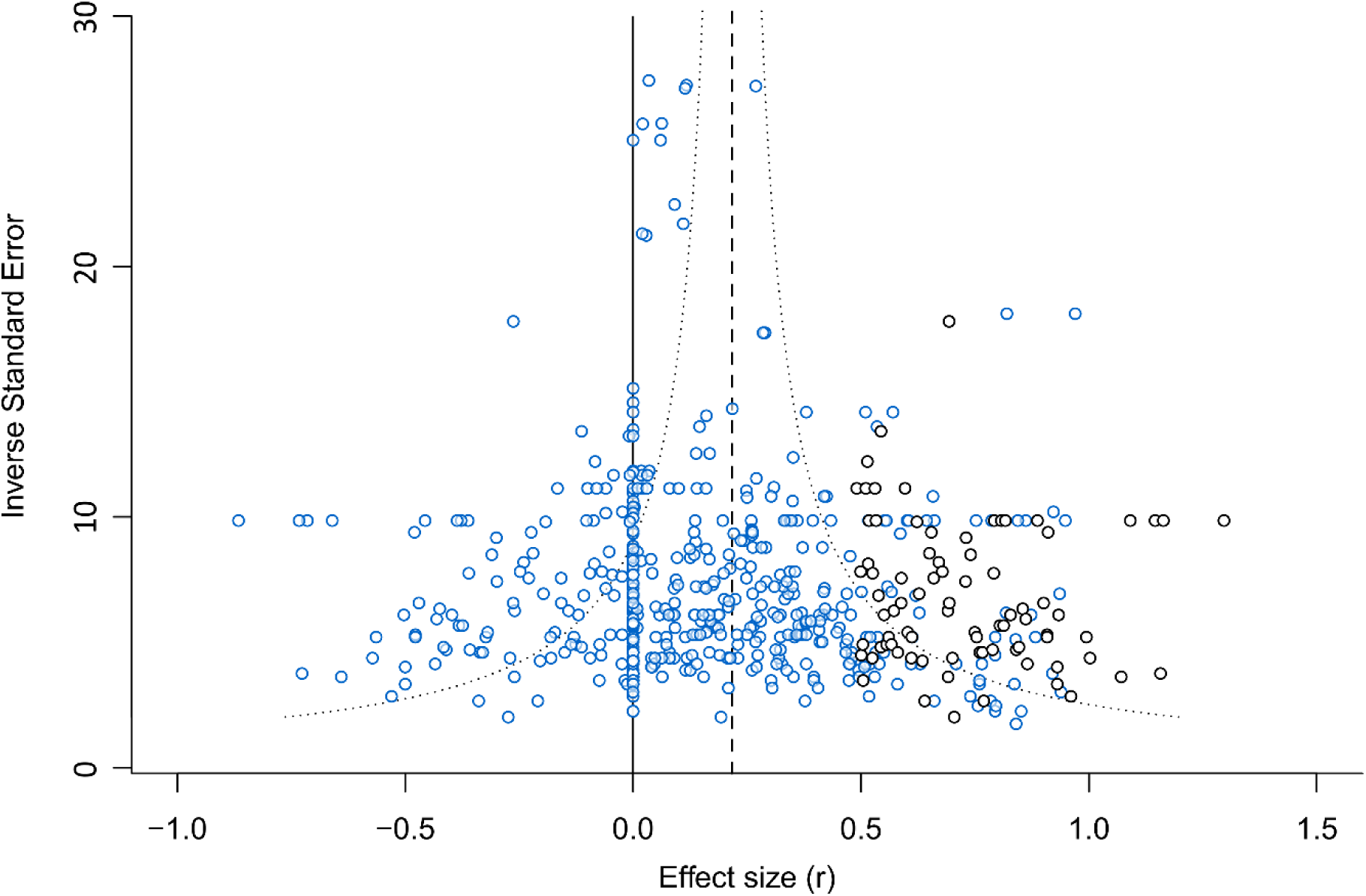
Funnel plot showing the relationship between effect size (the correlation coefficient r) and inverse standard error after performing a trim-and-fill analysis. Observed effect sizes are shown in blue (k= 520) and the imputed ‘missing’ effect sizes in black (k= 82). The dashed line shows the overall mean effect size estimate from a random-effects meta-analysis model after imputed the 82 ‘missing’ effect sizes. The dotted line illustrates the typical expected ‘funnel’ shape, with large sample-size studies resulting in effect size estimates closer to the mean.

